# Evolution and development of segmented body plan revealed by *engrailed* and *wnt1* gene expression in the annelid *Alitta virens*

**DOI:** 10.1101/2025.01.26.634940

**Authors:** Arsenii I. Kairov, Vitaly V. Kozin

## Abstract

Segmentation is one of the most striking features of bilaterians, and understanding its mechanisms provides insights into the evolution of body plans. In annelids, segmentation occurs at different developmental stages through a variety of processes, yet the molecular pathways remain underexplored. Aiming to compare segmentation patterns in ontogeny and phylogeny, we analysed the expression of *Avi-en* (homologous to *engrailed*) and *Avi-wnt1* in the nereidid polychaete *Alitta virens*. Using in situ hybridization, immunofluorescence, and cell proliferation assays, we mapped the spatiotemporal expression of these genes across embryonic, larval, and postlarval stages. We found that *Avi-en* was expressed in solid lateral domains early in the unsegmented protrochophore stage and progressed through a metameric pattern, while *Avi-wnt1* expression appeared later, also aligning with segmental boundaries. At the nectochaete stage, the posterior domain of *Avi-en* expression in the growth zone expanded and split into two due to increased cell proliferation. The postlarval segment primordium then developed progressively, culminating in the activation of *Avi-wnt1* at the posterior border. According to available published data, the revealed pattern of gradual segment formation is unique to nereidids. The observed divergence in gene expression and cell proliferation across annelids suggests that segmentation in bilaterians did not arise from a common ancestral mechanism. Our study enhances future progress in understanding the evolution of body patterning by providing a foundation for future comparisons.

## Introduction

The segmentation of the annelid body has been a central focus of evolutionary developmental biology for many years (Irvine and Martindale, 1996; Shimizu and Nakamoto, 2001; Balavoine, 2014; Zattara and Weisblat, 2020). Annelids exhibit a wide variety of anatomical patterns, including homonomously segmented errant polychaetes (which have more or less identical metameres), heteronomously segmented sedentary worms (whose metameres are functionally specialized and form different tagmata), and unsegmented sipunculids and echiurids. Segment formation occurs at different stages of ontogenesis, depending on the species: throughout life, as in oligochaetes; only in postembryonic development, as in most polychaetes; exclusively during the larval period, as in Pectinariidae; or only during embryogenesis, as in leeches. This results in varying number of segments, from oligomeric to polymeric species. The mechanisms responsible for dividing tissue layers into metameric units are equally diverse (Minelli and Fusco, 2004; Hannibal and Patel, 2013; Simsek and Özbudak, 2023). In different phylogenetic clades they range from boundary-driven to lineage-driven processes and involve different types of growth zones.

In polychaetes, the larval ectoderm of a solid trunk region (the somatic plate) is segmented through the formation of metameric ciliary rings and intersegmental furrows, which also subdivide mesodermal tissue (Anderson, 1973). Although this boundary-driven segmentation is considered characteristic of polychaetes, some species (*Platynereis, Scoloplo*s) show mesodermal bands with a teloblastic origin, suggesting clonal segmental domains (Fischer and Arendt, 2013; Özpolat et al., 2017). In contrast, in clitellate embryos, segmental mesoderm and ectoderm arise from a teloblastic growth zone. Following their budding off from teloblasts, the ectodermal and mesodermal daughter cells divide in a stereotypical manner, intermix, and generate anatomical segments, often with a shift in register relative to the clonal domains (Weisblat and Winchell, 2020). This embryonic pattern of clitellates and some crustaceans is referred to as lineage-driven segmentation. However, teloblasts do not persist beyond embryogenesis, and the growth zone organization in adult oligochaetes remains unclear. Non-clitellate annelids (i.e., “polychaetes”) also possess a growth zone, but it lacks identifiable teloblasts and begins producing segments only later in larval life. The precise nature of the growth zone in polychaetes, whether teloblastic or diffuse, remains uncertain (Kairov and Kozin, 2023). Discrepancies in the formation of larval segments from the trochophore’s hyposphere and postlarval segments from the growth zone suggest different developmental mechanisms and evolutionary implications. This is reflected in the ongoing debate surrounding the theory of primary heteronomy of segments (Iwanoff, 1928; Giangrande and Gambi, 1998). Thus, the diversity of developmental and anatomical segmentation patterns in annelids underscores their critical importance for comparative evolutionary analysis.

The molecular and cellular basis of segmentation in annelids is still not fully understood. The one clear consensus in the study of annelid segment formation is the critical role of cell proliferation, which drives axial elongation (de Rosa et al., 2005; Seaver et al., 2005; Paulus and Müller, 2006; Zattara and Bely, 2013; Ribeiro et al., 2021; Shalaeva and Kozin, 2023). Several genes are known to exhibit metameric expression patterns, including Wnt ligands, *engrailed*, *hes/hey*, *twist*, NK-genes, and members of the Hedgehog signaling pathway (Seaver et al., 2001; Prud’homme et al., 2003; Seaver and Kaneshige, 2006; Saudemont et al., 2008; Thamm and Seaver, 2008; Dray et al., 2010; Steinmetz et al., 2011; Gazave et al., 2014; Bastin et al., 2015; Kozin et al., 2016, 2019; Kairov and Kozin, 2023). Functional data are available for Wnt and Hedgehog signaling, which are essential for proper segmentation (Dray et al., 2010; Niwa et al., 2013). Less is known about the other genes, whether their metameric expression patterns are a cause of segmentation or merely a consequence of other patterning systems.

A particularly important segment patterning mechanism likely involves the Wnt signaling pathway. Wnt signaling is considered a conserved feature of bilaterians, playing a central role in anterior-posterior patterning, with activity concentrated at the posterior end (Martin and Kimelman, 2009; Petersen and Reddien, 2009). This is supported for annelids both by Wnt expression patterns and functional data (Dray et al., 2010; Pruitt et al., 2014; Bastin et al., 2015; Kozin et al., 2019; Ribeiro and Aguado, 2021). Furthermore, Wnt signaling regulates cell proliferation in the posterior growth zones of various bilaterians (Martin and Kimelman, 2009; Bénazéraf and Pourquié, 2013; Williams and Nagy, 2017; Constantinou et al., 2020; Mundaca-Escobar et al., 2022), which is especially relevant in the context of segmentation. Given this central role of Wnt pathway, a candidate gene approach focusing on Wnt associated positional markers offers a powerful strategy to dissect the mechanisms of segmentation.

Homologs of the segment polarity genes *engrailed* and *wingless* have an evolutionary conserved role in morphogenetic field delimitation. While their expression has been partially documented in nereidids (Prud’homme et al., 2003; Steinmetz et al., 2011; Kairov and Kozin, 2023) and other polychaetes (Seaver et al., 2001; Seaver and Kaneshige, 2006), no comprehensive analysis has been conducted to date. Specifically, previous studies have not explored continuous gene expression during larval and postlarval development, nor have they systematically compared the expression patterns of the orthologous genes. Here we examined the coordinated expression of segmental boundary markers *Avi-en* and *Avi-wnt1* throughout the developmental stages of *Alitta virens* with the goal of identifying shared mechanistic principles underlying segment formation in annelids.

*A. virens* is a nereidid polychaete whose embryonic and larval development (Dondua, 1975; Sveshnikov, 1978; Kairov and Kozin, 2023) closely resembles that of *Platynereis dumerilii* (Fischer et al., 2010). During gradual metamorphosis of nereidids (Wilson, 1892; Sveshnikov, 1978; Fischer et al., 2010), the trochophore transforms into a metameric larva. The hyposphere of the metatrochophore elongates via convergent extension, and the pygidial lobes separate from the trunk. The first morphological signs of metameric organization include additional ciliary bands called paratrochs, parapodia, and segmental grooves. The metatrochophore of nereidids develops four segments: an anterior underdeveloped cryptic (zero) segment and three chaetigerous segments. Following the emergence of head appendages and initiation of parapodia movements, the larva transforms into a nectochaete. The nectochaete remains planktonic before eventually settling on the substrate. This process aligns with the onset of feeding, which occurs asynchronously among individuals. Subsequent postlarval segments are formed through anamorphic growth, involving intercalation in front of the pygidium. The well-described and accessible developmental stages of nereidid polychaetes make them a valuable model for investigating the mechanisms of segment addition.

In this study, we employed a combination of in situ hybridization, immunofluorescence, and cell proliferation assays to map the spatiotemporal expression of the segment polarity genes *Avi-en* and *Avi-wnt1* throughout the embryonic, larval, and postlarval development of the annelid *A. virens*. By characterizing the dynamic patterns of these markers across the developmental stages, we provide a comprehensive overview of the molecular events underlying segment formation in this nereidid polychaete. Our data provide a necessary foundation for future comparative studies on the evolution of segmentation in annelids and beyond.

## Materials and Methods

### Animals and fixation

Adult *A. virens* worms were collected at the White Sea near the Marine Biological Station of St. Petersburg State University. The artificial fertilization and embryo culturing were performed as described earlier (Dondua, 1975). Embryos and early larval stages were cultivated at +14°С, then temperature was increased to +20°C from the early nectochaete stage. Larvae and juveniles were fixed in 4% paraformaldehyde on 1xPBS/0.1%Tween20 overnight at +4°C, followed by washes in 1xPBS/0.1%Tween20 and dehydrated in 100% methanole. Starting from mid metatrochophore stage (135 h.p.f.), the larvae were anaesthetised in MgCl2 before fixation.

### EdU labeling

To reveal cell proliferation, larvae and juveniles were incubated in 50 μM EdU (5-ethynyl-2′-deoxyuridine) for 15 minutes before anesthetization and fixation. Click-reaction using sulfo-Cyanine5 azide (Lumiprobe) was performed as described earlier (Shalaeva and Kozin, 2023).

### Whole mount *in situ* hybridization (WMISH) and immunohistochemistry

Phylogenetic assignment of the genes of interest and their expression patterns at some stages in nereidid polychaetes have been reported previously (Prud’homme et al., 2003; Janssen et al., 2010; Steinmetz et al., 2011; Kozin et al., 2019; Kairov and Kozin, 2023). Plasmids with cloned fragments of *Avi-en* and *Avi-wnt1* (∼1,6 kb) were provided by R. P. Kostyuchenko. Linearized plasmids were used for synthesis of DIG-labelled RNA probes. From 50 to 100 specimens of each developmental stage were analysed using whole mount in situ hybridization according to the protocol described earlier (Shalaeva et al., 2021). Incubation with RNA probes lasted 48 h; staining with BCIP/NBT lasted from 20 to 40 h. For simultaneous detection of two genes, RNA probes were mixed at equimolar concentrations.

### To combine WMISH and EdU detection we performed standard WMISH protocol followed by standard click-reaction

For combined WMISH and antibody labeling we used the same WMISH protocol with following modifications: samples were incubated simultaneously with anti-DIG antibodies and primary mouse anti-acetylated tubulin antibody (Sigma T6793, dilution 1:400). Antibody washing was followed by standard NBT/BCIP staining. Stained samples were washed in 1xPBS/0.1%Tween20 and incubated in 5% sheep serum for 1 hour followed by incubation with secondary antibodies (Sigma T5168 anti-mouse antibodies conjugated with Alexa-488 fluorescent dye, dilution 1:500) overnight at +4 °C. The washed samples were mounted in 90% glycerol.

### Data Visualization

Confocal images were obtained using Leica TCS SPE confocal microscope. WMISH results were visualized using DIC optics with Axio Imager D1 microscope (Carl Zeiss). Confocal detection of the WMISH signal was performed as described earlier (Jekely and Arendt, 2007). All schemes and figures were made with Adobe Illustrator, Adobe Photoshop and ImageJ.

## Results

### *Avi-en* expression

*Avi-en* mRNA was first detected at 32 hours post-fertilization (h.p.f.) during the protrochophore stage. Transcripts were localized in five groups of cells arranged in a bilaterally symmetrical pattern (Fig. 1A, B). A medial, unpaired domain was observed near the vegetal pole (Fig. 1A, B, black arrow). Additionally, two pairs of bilaterally symmetrical domains were located on the lateral sides of the hyposphere, with the anterior domains being larger than the others.

**Fig. 1.**
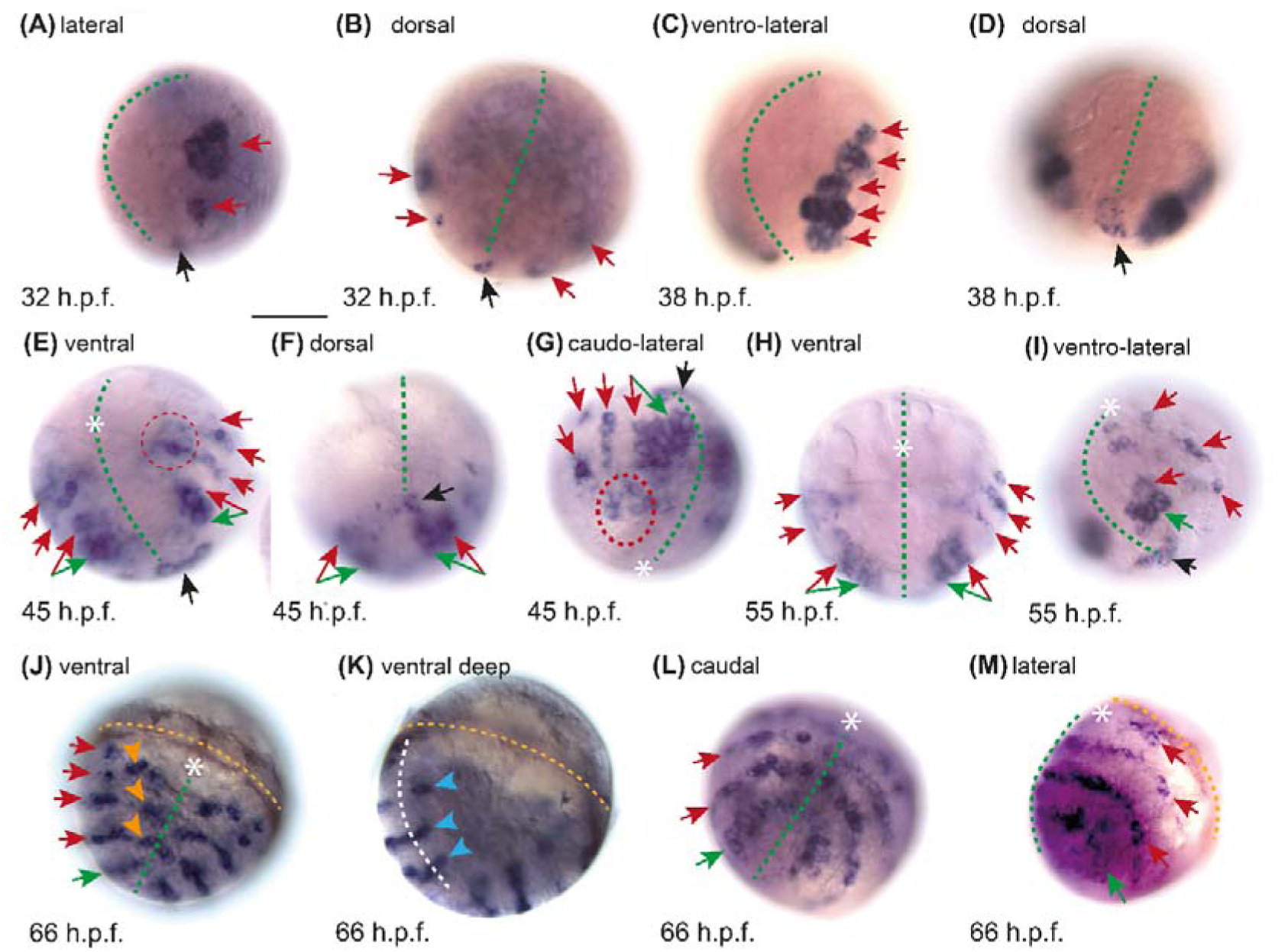
*Avi-en* expression during early larval stages in *A. virens*. Red arrows — metameric domains, black arrow — unpaired domain, green arrow — expression in the presumptive growth zone, orange arrowheads — expression in the neuroectodermal cells, white asterisk — stomodeum, yellow dotted line — prototroch, green dotted line — medial line of the body, red dotted line — area of connection between metameric domains. Scale bar — 50 μm. (A) Protrochophore, 32 hours post fertilization (h.p.f.), lateral view, animal pole is to the top, ventral side is to the left. (B) Protrochophore, 32 h.p.f., dorsal view, deep focus. (C) Protrochophore, 38 h.p.f., lateral view. (D) Protrochophore, 38 h.p.f., ventro-caudal view. (E) Protrochophore, 45 h.p.f., ventral view, anterior is to the top. (F) Protrochophore, 45 h.p.f., dorso-caudal view, anterior is to the top. (G) Protrochophore, 45 h.p.f., caudal-lateral view, ventral side is to the top. (H) Early trochophore, 55 h.p.f., ventral view, anterior is to the top, deep focus. (I) Early trochophore, 55 h.p.f., lateral view, ventral is to the left top corner. (J) Mid-trochophore, 66 h.p.f., ventral view, anterior is to the top. (K) The same object as in (J), deep focus. White dotted line — border between ectoderm and mesoderm. Blue arrowheads — mesodermal domains. (L) Mid-trochophore, 66 h.p.f., ventro-caudal view, anterior is to the top. (M) Mid-trochophore, 66 h.p.f., lateral view, anterior is to the top right corner.

By 38 h.p.f., *Avi-en* mRNA expression was evident in rows of cells along the lateral sides, stretching in the anterior-posterior direction (Fig. 1C). These rows were organized into five groups of cells, which later align with the segment borders. The unpaired domain expanded in size and connected posteriorly with the lateral domains on the dorsal side (Fig. 1D, black arrow).

At 45 h.p.f., the expression pattern consisted of four pairs of bands arranged perpendicular to the anterior-posterior axis, located on the lateral sides of the body (Fig. 1E). These bands corresponded to the distinct cell groups observed in the previous stage. The most anterior band was the shortest, while the other three bands were longer and interconnected at their ventrolateral ends (Fig. 1E, G). The last pair of bands was the widest and corresponded to the two undiverged domains observed in the 38 h.p.f. embryo (Fig. 1E, F, G, red and green arrows). The posterior unpaired domain connected these wide posterior bands on the dorsal side (Fig. 1F, black arrow).

At the early trochophore stage (55 h.p.f.), the *Avi-en* expression pattern consists of four pairs of bands (Fig. 1H, I). Unlike the previous stage, these bands no longer connect at the ventrolateral sides (Fig. 1I). The staining within the posteriormost domain appears discontinuous, with a medial area of weaker or absent signal (Fig. 1H), which may indicate the initial event leading to the separation of this domain into distinct parts, as is clearly observed at later stages. The posterior unpaired domain persists on the dorsal side (Fig. 1I, black arrow).

By the middle trochophore stage (66 h.p.f.), *Avi-en* mRNA was detected in five pairs of stripes of ectodermal cells arranged in a bilaterally symmetrical pattern (Fig. 1J, M). A corresponding metameric signal was observed in internal (mesodermal) cells, aligning with the superficial ectodermal domains (Fig. 1K). The most anterior pair of stripes was the shortest and restricted to the ventrolateral regions of the larva. The remaining stripes were curved in an arc, with the left and right halves joining at the ventral side but not extending to the dorsal midline. Cells located at the ventral side, which are neuroectodermal in nature, exhibited a posterior phase shift relative to the lateral regions of expression, indicating a mosaic pattern. The most posterior domain formed an almost circular shape, corresponding to the prospective growth zone (Fig. 1L).

At the early metatrochophore stage (116 h.p.f.), the *Avi-en* expression pattern remained largely unchanged. There were five metameric transverse bands, which were circular except for the most anterior one, which was interrupted on the ventral and dorsal sides (Fig. 2A, B). The expression pattern became more mosaic, as the lateral domain parts were now distinct from the ventral (neuroectodermal) regions. Metatrochophores exhibited external segmentation features, including segmental grooves and paratrochs (metameric circular ciliary bands), enabling us to correlate metameric domains with segment borders. The first and second stripes were located between the prototroch and the first intersegmental groove. The third and fourth expression domains aligned with segmental grooves at the boundaries between segments 1/2 and 2/3, respectively. The posterior-most expression band corresponded to the border between the last segment and the area of the future pygidium (Fig. 2B, green arrow).

**Fig. 2.**
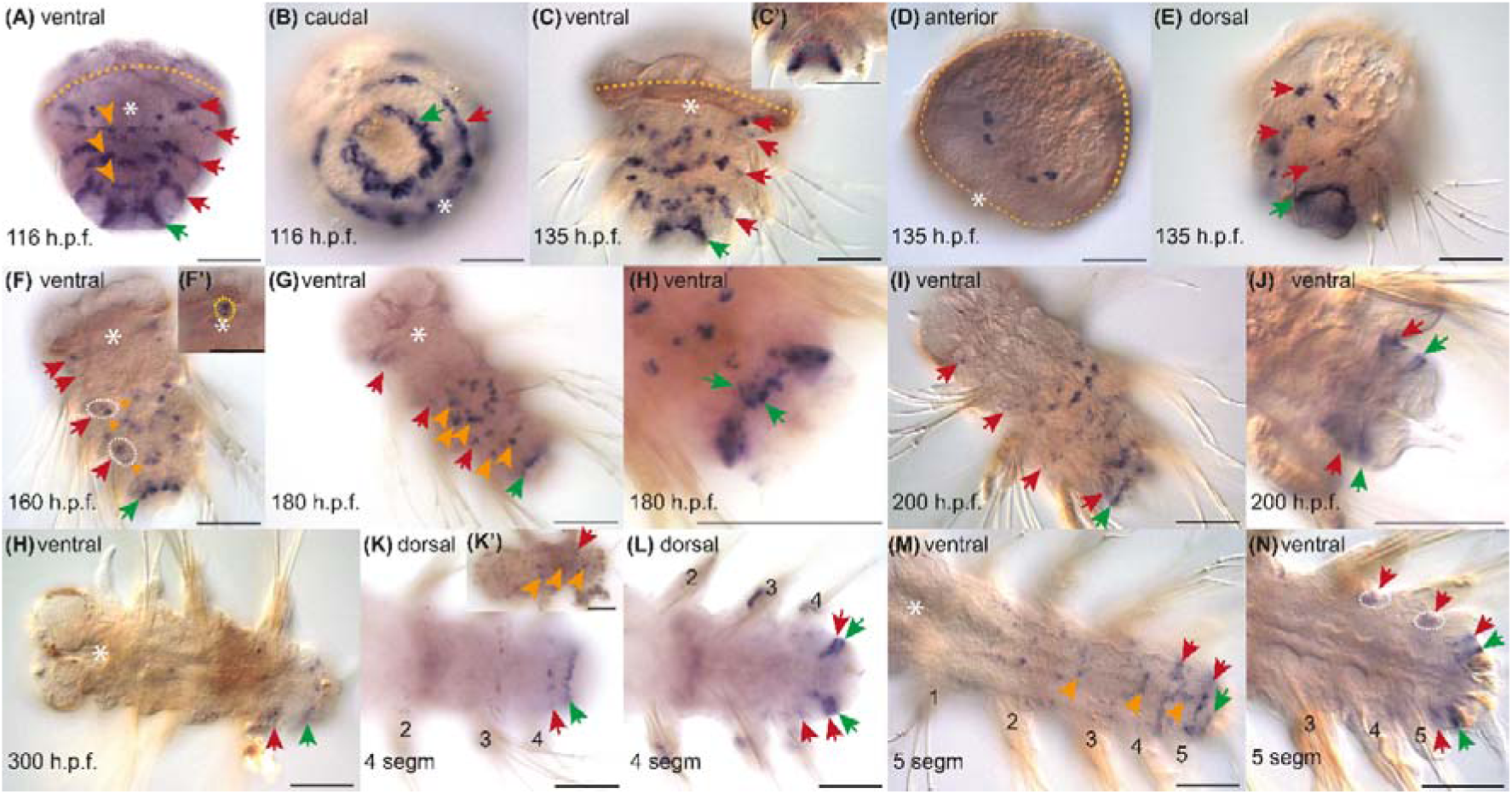
*Avi-en* expression during segmented larvae and juvenile stages in *A. virens*. Scale bars — 50 μm. Red arrows — metameric domains, green arrow — expression in the presumptive growth zone, orange arrowhead — expression in the neuroectodermal cells, yellow dotted line — prototroch, white asteriks — stomodeum or pharynx. White dotted lines emphasize metameric domains extended on two adjacent segments. (A), (C), (F), (F’) — ventral view, anterior is to the top; (B) — caudal view, ventral is to the right bottom corner; (G), (H), (I), (J) — ventral views, anterior is to the top left corner; (D) — anterior view; (H), (K’), (M), (N) — ventral views, anterior is to the left; (K), (L) — dorsal views, anterior is to the left. (A) Early metatrochophore, 116 h.p.f. Expression is similar to that of the mid trochophore. (B) Early metatrochophore, 116 h.p.f. *Avi-en*+ cell bands are circumferential. (C) Mid metatrochophore, 135 h.p.f. (C’) *Avi-en*+ mesodermal cells (red dotted line) in the area of the future growth zone. (D) Mid-metatrochophore, 135 h.p.f., episphere contains 4 *Avi-en*+ groups of cells closer to the dorsal side. (E) Mid-metatrochophore, dorso-lateral view, anterior is to the top. Metameric domains expand to the ventral and dorsal sides. (F) Late metatrochophore, 160 h.p.f. (F’) Magnified image of the pharynx. Some specimens have *Avi-en*+ cells (yellow dotted line) in the anterior pharynx. (G) Early nectochaete, 180 h.p.f. (H) Early nectochaete, the same object as in (G), magnified image of caudal end, deep focus. There are two adjacent rows of *Avi-en*+ cells in the growth zone (green arrowheads). (I) Mid-nectochaete, 200 h.p.f. (J) Mid-nectochaete, the same object as in (I), magnified image of the caudal end, deep focus. There are two bands of *Avi-en*+ cells in the pygidium. (H) Juvenile with the forming 4^th^ chaetigerous (1^st^ postlarval) segment, 300 h.p.f. The expression signal is absent in neuroectodermal cells. (K)-(L) Juvenile with the 4^th^ segment. Two rings of *Avi-en*+ cells in the pygidium. (K’) Expression signal is observed in neuroectodermal cells in larval segments (orange arrowheads). (L) The same object as in (K), deep focus. Two rows of *Avi-en*+ cells indicate the formation of a new segment. (M), (N) 5-segment juvenile. Each segment is numbered with an Arabic numeral.

In nereidid polychaetes, paratrochs are situated at the posterior part of each segment (Fischer et al., 2010; Bastin et al., 2023), and the growth zone lies between the third paratroch and the telotroch (Nielsen, 2004; Fischer et al., 2010; Starunov et al., 2015). The relative positions of *engrailed* expression stripes and ciliary bands have been previously documented only in *P. dumerilii* (Steinmetz et al., 2011). Using confocal microscopy, we observed that the third and fourth metameric *Avi-en* domains were positioned posterior to, and did not abut, the paratrochs (Fig. 3A), corresponding to the anterior parts of chaetigerous segments 2 and 3. The posterior-most circular domain was located between the last paratroch and the telotroch, indicating that it lies at the anterior border of the pygidium and coincides with the prospective growth zone. The two anterior expression stripes were not confined to morphological boundaries. The second expression domain was located posterior to the metatroch, while the first domain was anterior to it (Fig. 3A, B). Consequently, the first and second domains correspond to the anterior regions of segment 0 (a cryptic segment) and segment 1, respectively.

**Fig. 3.**
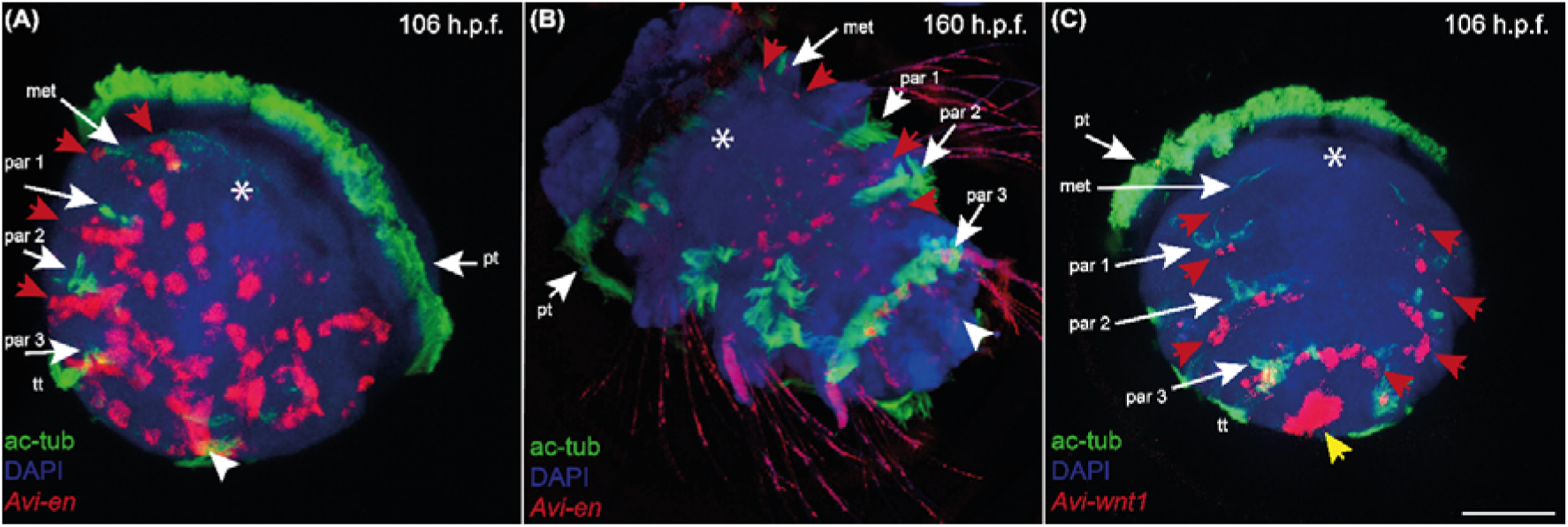
*Avi-en* and *Avi-wnt1* expression (red channel) relative to ciliary bands (green channel). Maximum projections of confocal Z-stacks of ventral views, anterior is to the top. Scale bar — 50 μm. White asterisk — stomodeum/pharynx. Par 1…3 — paratrochs of the 1^st^ - 3^rd^ segments, pt — prototroch, tt — telotroch, met — metatroch, red arrows — metameric expression domains, white arrowhead — expression domain in the growth zone, yellow arrow — terminal expression domain. (A) *Avi-en* expression, early metatrochophore. 3^rd^ and 4^th^ rows of expression are posterior to the 1^st^ and 2^nd^ paratrochs, 5^th^ row is located between the 3^rd^ paratroch and telotroch. (B) *Avi-en* expression, late metatrochophore. (C) *Avi-wnt1* expression, early metatrochophore. Metameric domains are located posterior to metatroch/paratrochs. The most posterior band of expression is located between the 3^rd^ paratroch and telotroch.

At the mid-metatrochophore stage (135 h.p.f.), the expression pattern remains largely unchanged from the previous stage and consists of five metameric domains. The last two domains are circular, while the others are interrupted on the dorsal side of the body (Fig. 2C, E). Ventromedial neuroectodermal *Avi-en*-positive cells increase in number and shift further posteriorly relative to the flanking lateral fragments. This shift enhances the mosaic pattern, as neuroectodermal cells no longer form distinct bands with the ventrolateral domains (Fig. 2C). Within the last expression domain, which corresponds to the prospective growth zone, *Avi-en* mRNA was detected in ectodermal and also in deep mesodermal cells (Fig. 2C’, red dotted line). Additionally, *Avi-en* expression was observed in four groups of cells located in the episphere, closer to the dorsal side (Fig. 2D).

At the late metatrochophore stage (160 h.p.f.), the *Avi-en* expression pattern becomes increasingly mosaic (Fig. 2F, 3B). Unlike the stable circular posterior domain, ventral neuroectodermal cells no longer form distinguishable bands with the ventrolateral ectodermal cells. However, the ventrolateral domains can still be grouped into four pairs. *Avi-en* mRNA disappeared from the lateral and dorsal regions of the larval body. The third and fourth metameric domains span both sides of the furrows, extending across the boundaries of two adjacent segments (Fig. 2F, white dotted line). Additionally, some specimens demonstrated *Avi-en*-positive cells in the pharynx (Fig. 2F’).

Previously, we described the *Avi-en* expression pattern during the final stages of metamorphosis in *A. virens* (Kairov and Kozin, 2023). In this study, we confirm our earlier conclusions through additional analysis using morphological and proliferative markers. At the early nectochaete stage (180 h.p.f.), the most significant changes were observed in the posterior ring-shaped domain at the boundary between the third segment and the pygidium (Fig. 2G). This domain expanded along the anterior-posterior axis, forming two rows of cells instead of one, as seen in earlier stages. The expansion occurred progressively from the ventral to the dorsal side (Fig. 2H). Anteriorly, the signal disappeared from the episphere.

At 200 h.p.f., nectochaetes exhibited two circular rows of *Avi-en*-positive cells in the anterior part of the pygidium (Fig. 2I, J). These rows likely represent two parts of *Avi-en* expression domain that were adjacent at the early nectochaete stage but diverged during cell proliferation and pygidium growth.

To identify exact areas of cell proliferation, we performed an EdU assay. At 200 h.p.f., EdU incorporation was detected in the anterior end of the nectochaete (episphere), in the trunk neuroectoderm, and prominently in the pygidium (Fig. 4A). In the pygidium, EdU labeling was especially intensive, with a lower density of labelled nuclei in its posterior region. During the formation of the fourth segment (Fig. 4B, C), the distribution of EdU-labelled nuclei resembled that in nectochaetes. However, after the separation of the new segment, it showed the highest labeling density, whereas the pygidium contained fewer EdU-labeled nuclei. This confirms that the pygidium and the emerging segment are regions of active cell proliferation.

**Fig. 4.**
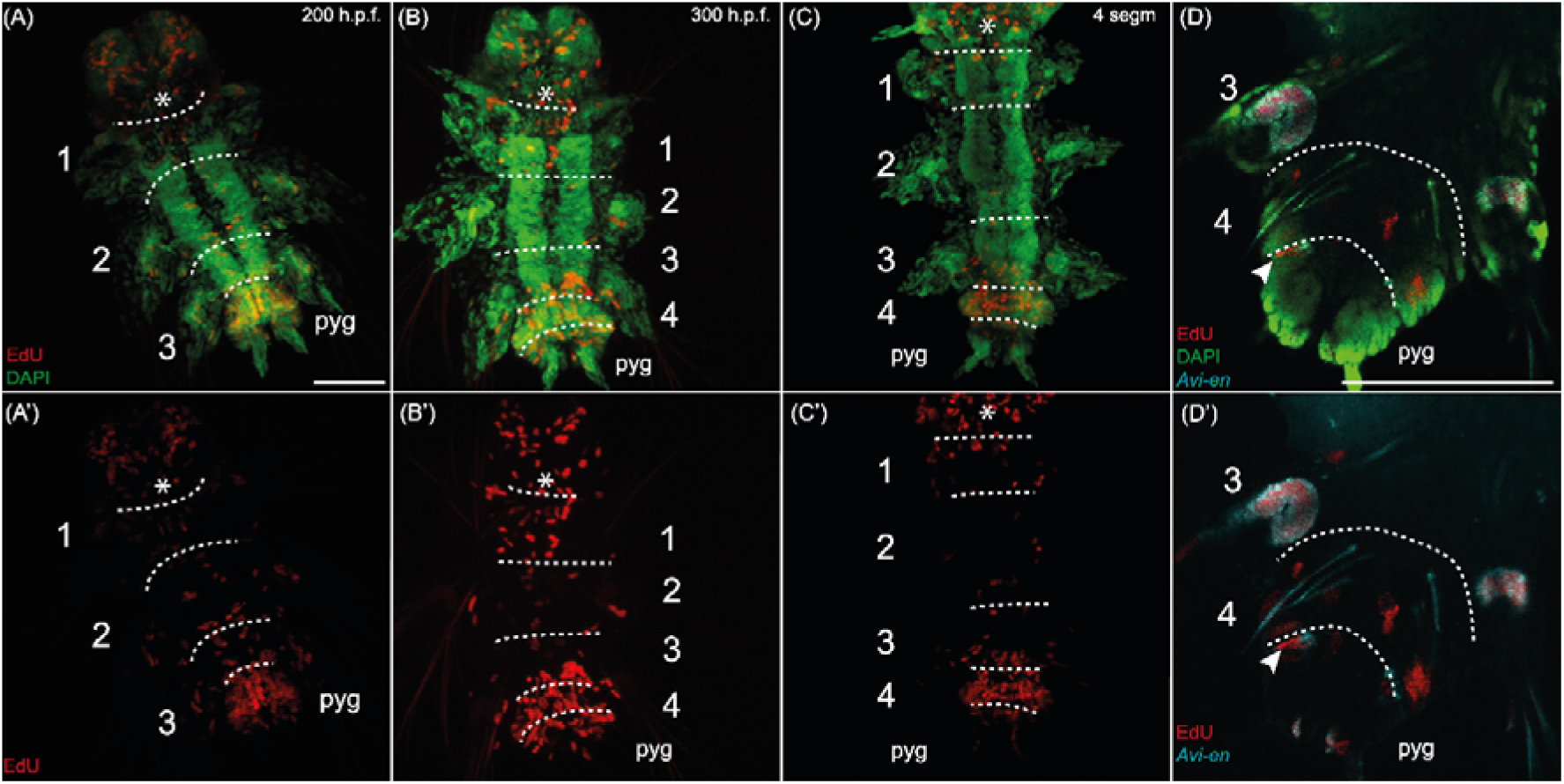
Cell proliferation reviled by EdU incorporation (red channel) during the first postlarval segment development in *A. virens* nectochaetes and juveniles. Nuclei were stained with DAPI (green channel). (A)-(C) Maximum projections of confocal Z-stack of ventral views, (D) Separate confocal section. Anterior is to the top. Scale bar — 50 μm. White asterisk — pharynx, white dotted line — intersegmental borders, pyg — pygidium. Each segment is numbered with an Arabic numeral. (A) Mid-nectochaete, 200 h.p.f. (B) Juvenile with the forming 4^th^ chaetigerous (1^st^ postlarval) segment, 300 h.p.f. (C-D) 4-segment juvenile. (D) EdU assay combined with *Avi-en in situ* hybridization. *Avi-en*+ cells (cyan channel) colocalize with the EdU+ nuclei in the growth zone (white arrowhead).

Following the appearance of a segmental groove within the pygidium, specifically during the growth and morphogenesis of the fourth segment, the signal was no longer detected in the larval segments (Fig. 2H). Towards the posterior part of the body, two *Avi-en* expression domains were observed: one located at the anterior border of the developing fourth segment (the first postlarval segment), and the other along the anterior border of the pygidium, corresponding to the growth zone of juvenile worms. Once the first postlarval segment is fully formed, expression in neuroectodermal cells resumes anterior to the fourth segment (Fig. 2H’). *Avi-en*-positive cells were also detected at the anterior border of the fourth segment and the anterior border of the pygidium (Fig. 2K). In more mature individuals with elongated pygidium, two bands of cells were observed within the pygidium, similar to those seen in the middle nectochaete stage (Fig. 2L).

The expression pattern of *Avi-en* in five-segment juveniles was found to be comparable to that observed in four-segment juveniles (Fig. 2M). The signal was detected in neural cells and at the boundaries between posterior segments. Within the pygidium, two circular rows of cells exhibiting the signal were retained: one in a more anterior position (at the anterior morphological boundary of the pygidium) and another located posterior to the first one (Fig. 2N). The distance between these two rings varied among individuals, reflecting the ongoing growth process of the segment primordium. In fully differentiated postlarval segments (the fourth and fifth chaetigers), metameric *Avi-en* expression domains were positioned on both sides of the groove, spanning the territory of two adjacent segments (Fig. 2N, white dotted line).

### *Avi-wnt1* expression

Previously, *Avi-wnt1* expression was partially described in *A. virens* trochophores (Kozin et al., 2019). In the present study, we detected the signal at an earlier stage than previously reported and conducted a detailed analysis of segmented larvae and juveniles. At 38 h.p.f., *Avi-wnt1* transcripts were localized in superficial cells at the posterior pole of the embryo (Fig. 5A). In 45 h.p.f. protrochophores, the posterior expression domain slightly expanded, but no new domains appeared (Fig. 5B).

**Fig. 5.**
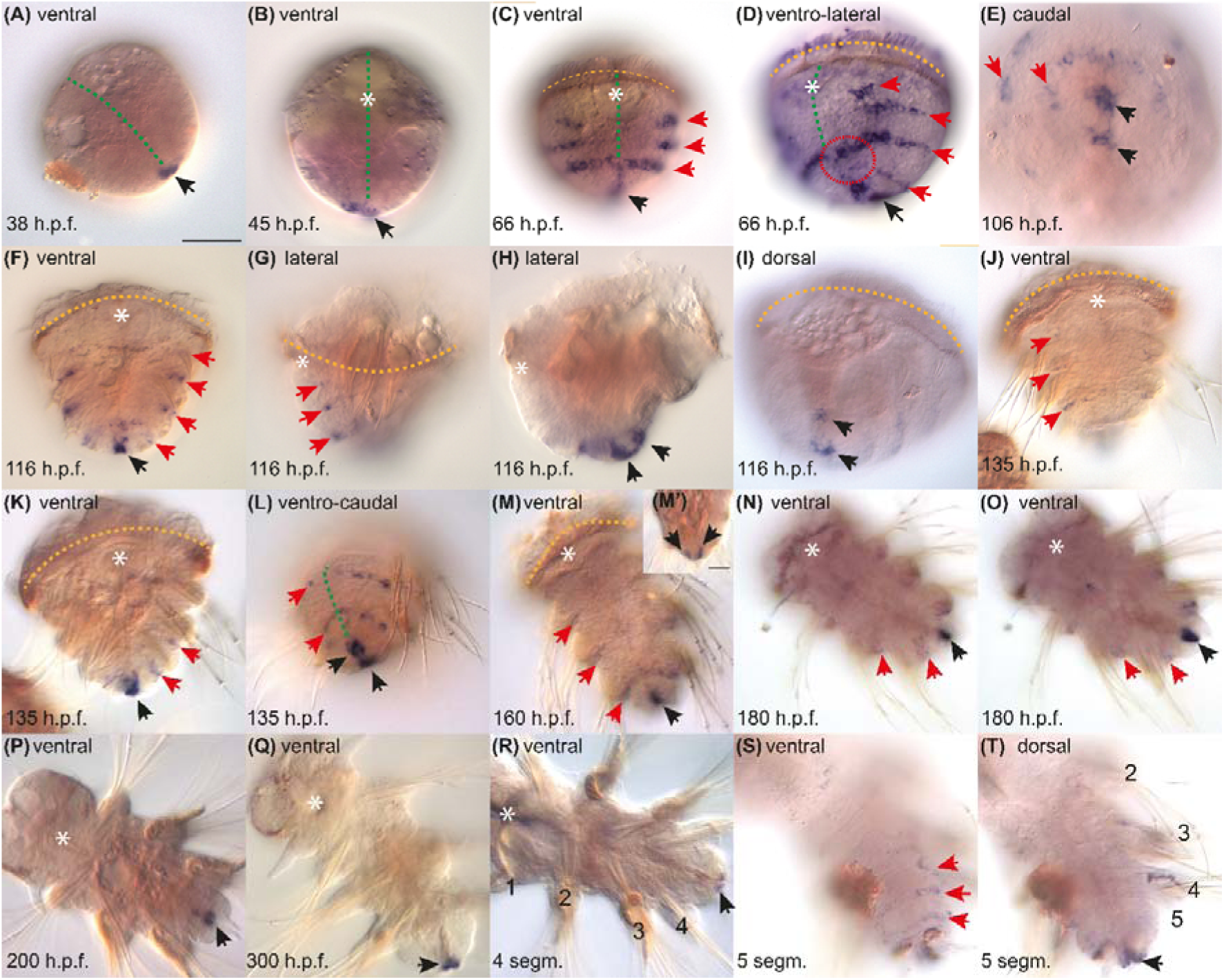
*Avi-wnt1* expression in *A. virens* development. Black arrow — caudal domain of expression, red arrows — metameric domains of expression. (A), (M), (N), (O), (P), (Q), (R) — ventral view, anterior is to the top left corner; (B), (C), (F), (J), (K) — ventral view, anterior is to the top; (D), (G), (H) — lateral view, anterior is to the top, ventral is to the left; (E) — caudal view, ventral is to the top. (I) — dorso-caudal view, anterior is to the top; (H) — ventro-caudal view, anterior is to the top; (S), (T) — dorsal view, anterior is to the top left corner. (A) Protrochophore, 38h.p.f. Only the terminal posterior domain of expression is detected. (B) Protrochophore, 45h.p.f. (C), (D) Mid-trochophore, 66 h.p.f. Metameric bands arise at ventro-lateral sides. Red dotted line — area of connection between metameric domains (E) Early metatrochophore, 106 h.p.f. (F)-(I) Early metatrochophore, 116 h.p.f. Metameric bands of expression are located along segmental borders. (H), (I) Terminal posterior domain of expression consists of two parts. (J)-(L) Mid-metatrochophore, 135 h.p.f. (M), (M’) Late metatrochophore, 160 h.p.f. (M’) Lateral view, deep focal plane, ventral is to the right. The terminal expression domain marks the hindgut. (N), (O) Early nectochaete, 180 h.p.f. (P) Mid-nectochaete, 200 h.p.f. Metameric domains are not detectable. (Q) Juvenile with the forming 4^th^ chaetigerous (1^st^ postlarval) segment, 300 h.p.f. (R) 4-segment juvenile. (S), (T) 5-segment juvenile. Metameric expression domains mark the borders of postlarval segments. Each segment is numbered with an Arabic numeral.

By the mid-trochophore stage (66 h.p.f.), the *Avi-wnt1* expression pattern became metameric. Four transverse bands of ectodermal cells expressing *Avi-wnt1* emerged on both lateral sides of the hyposphere (Fig. 5C). The two posterior metameric bands formed an arc and were continuous on the ventral side of the body (Fig. 5D, red dotted line). The posterior medial expression domain expanded anteriorly on the ventral side of the larval body and intersected with the last two metameric bands.

In metatrochophores, the *Avi-wnt1* expression pattern did not change significantly (Fig. 5E-F). The posterior expression domain in the proctodaeum region included two parts: a more ventral and a dorsal one (Fig. 5E, H, I). These domains incorporated both superficial and internal cells. The metameric expression bands extended along the segment grooves (Fig. 5F, G) and posterior to the paratrochs (Fig. 3C). Notably, these bands were positioned closer to the paratrochs than the *Avi-en* bands. The anterior-most *Avi-wnt1* expressing band was situated posterior to the metatroch (Fig. 3C).The most posterior metameric *Avi-wnt1* band was located in front of the telotroch closely opposed to the *Avi-en* expression ring in the future growth zone (Fig. 3C). To determine whether *Avi-en* and *Avi-wnt1* are co-expressed in the same cells or expressed in adjacent domains, we performed combined WMISH with a mixture of both probes. The resulting staining pattern revealed metameric bands that were broader than those observed for either gene alone (Fig. 6A, B). This arrangement indicates that *Avi-en* and *Avi-wnt1* are expressed in neighboring cell populations rather than being co-expressed within the same cells. Furthermore, combined probe staining revealed that the terminal posterior *Avi-wnt1* domain in the proctodaeum does not directly abut the most posterior *Avi-en* ring. Instead, the distinct internal *Avi-wnt1* signal is positioned beneath the more superficial posterior *Avi-en* domain, suggesting a potential signaling relationship where *wnt1*-expressing cells may influence the overlying *en*-expressing cells of the growth zone.

**Fig. 6.**
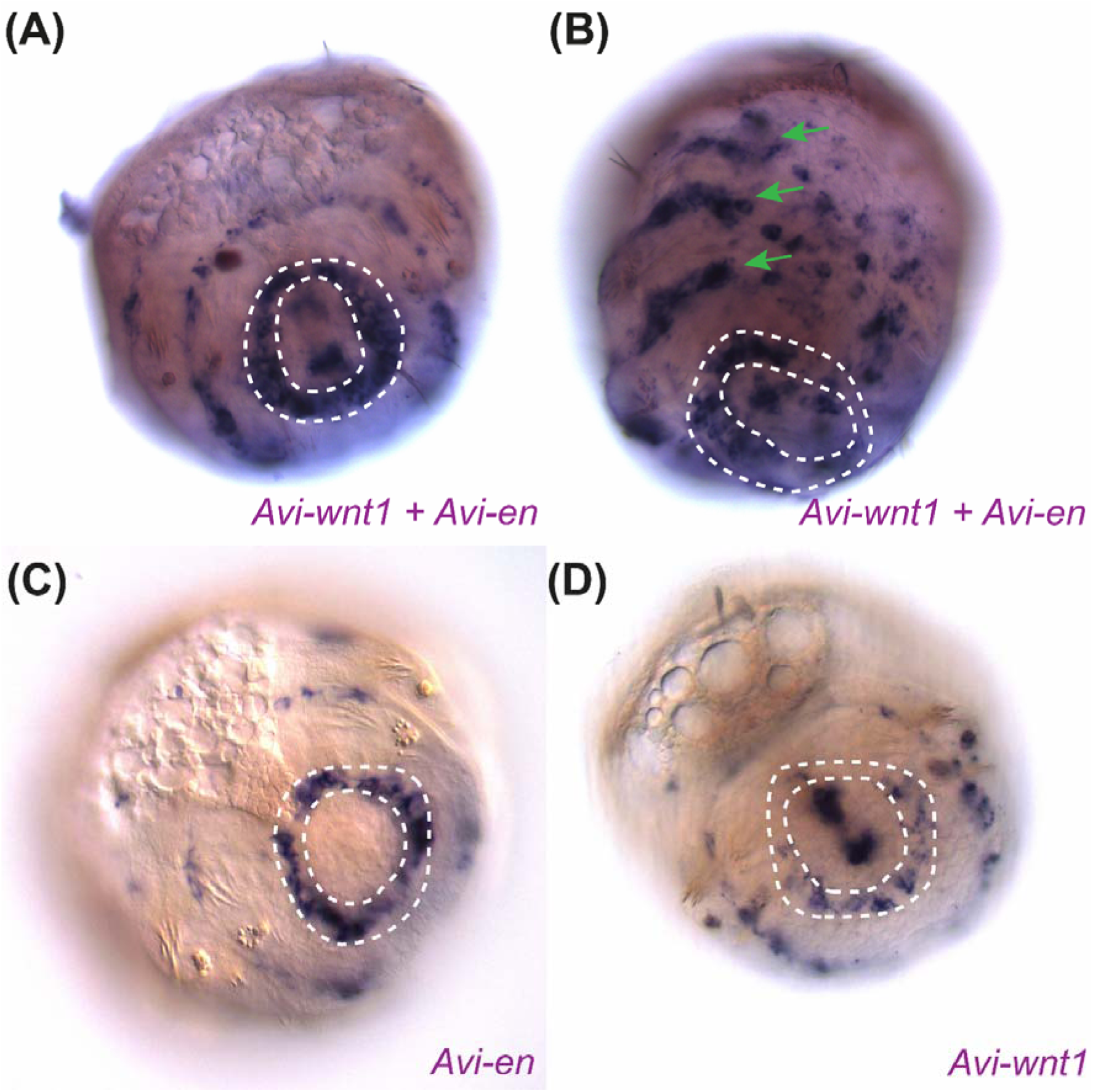
Comparative analysis of *Avi-en* and *Avi-wnt1* expression domains in the early metatrochophore (116 h.p.f.) suggests adjacent but largely non-overlapping expression. (A), (C), (D) Dorso-caudal views; dorsal is to the top left. (B) Ventro-lateral view; anterior is to the top, ventral is to the right. (A), (B) Whole-mount in situ hybridization with a mixture of *Avi-en* and *Avi-wnt1* DIG-labeled RNA probes. The staining domains are broader compared to individual gene expression. White dotted line indicates the boundary between the 3rd segment and the growth zone. Green arrows point to the metameric domains at the boundaries of segments 0, 1, 2, and 3, resulting from the combined staining of adjacent *Avi-en* and *Avi-wnt1* expression stripes. (C) Individual *Avi-en* expression pattern in a representative specimen. (D) Individual *Avi-wnt1* expression pattern in a representative specimen.

In middle and late metatrochophores, *Avi-wnt1* expression in metameric domains gradually faded and completely disappeared by the 200 h.p.f. nectochaete stage (Fig. 5J-P). However, in segmented larvae, the signal was detected in a circular band at the posterior border of the third segment (Fig. 5N, O). Terminal *Avi-wnt1* expression continued in the hindgut wall (on the ventral and dorsal sides) (Fig. 5M, M’, O, P).

At the onset of anamorphic growth and postlarval segment formation, the posterior medial *Avi-wnt1* expression domain decreased in length (Fig. 5Q, R). The signal was detected in the most terminal cells of the hindgut. By the time the sixth segment formed, the *Avi-wnt1* expression pattern became metameric again. In addition to the signal in the hindgut (Fig. 5T), mRNA was detected in cells at the posterior border of postlarval segments on the ventral and lateral sides of the body (Fig. 5S).

## Discussion

### Analysis of *Avi-en* and *Avi-wg* expression throughout the developmental stages of *A. virens*

*Avi-en* expression was first detected at the protrochophore stage. Initially, the expression was bilaterally symmetrical, with a single unpaired domain at the posterior end and two pairs of lateral domains (Fig. 6). These lateral domains gradually expanded along the anterior-posterior (AP) axis, forming two longitudinal bands subdivided into five distinct units. This expansion was likely driven by ongoing cell proliferation or the induction of *Avi-en* in new territories. In contrast, *Avi-wnt1* expression appeared later than *Avi-en*, suggesting that *Avi-wnt1* is not essential for the initiation of *Avi-en* expression. This timing is consistent with RNA-seq data from *Platynereis dumerilii* transcriptomic studies (Pruitt et al., 2014; Chou et al., 2016).

The five groups of *Avi-en*-expressing cells represent the future metameric domains, which became clearly visible by the mid-trochophore stage (Fig. 1, 7). These domains corresponded to the molecular boundaries of future body segments: the anterior four were confined to larval segments, while the posterior-most was associated with the pygidium and the prospective growth zone.

The formation of the *Avi-en* metameric pattern occurred progressively. Initially, domains corresponding to the cryptic (0), 1st, and 2nd segments became distinct. Later, the prospective pygidial and 3rd segmental domains became individualized. Examination of the domain shapes for segments 1 and 2 (Fig. 1E) revealed differences in the degree of separation between dorsal and ventral regions: while the ventral parts remained connected, the dorsal parts had already separated. Eventually, the posterior medial *Avi-en* domain merged with the last band associated with the pygidium and prospective growth zone. The divergence of *Avi-en*-positive cells was likely driven by cell migration from the dorsal to the lateral sides, a process previously observed during somatic plate formation (Wilson, 1892; Anderson, 1973). At the embryonic stages, the *Avi-en* signal was confined to the lateral sides, but by the mid-trochophore stage, its expression expanded ventrally to include the neuroectoderm. Rearrangements in neuroectodermal expression may have been caused by convergent extension within the neuroectoderm (Steinmetz et al., 2007).

*Avi-wnt1* expression was detected later, in cells at the posterior pole of protrochophores, with metameric domains appearing only in trochophores. During early larval development, the metameric bands of *Avi-wnt1* (except for the most posterior band) were positioned more anteriorly than the corresponding *Avi-en* bands. This aligns with known data showing that *wnt1* is expressed at the posterior border of body parts, while *engrailed* is expressed at the anterior border, as observed in *P. dumerilii* (Prud’homme et al., 2003) and other bilaterians (Janssen et al., 2010; Vellutini and Hejnol, 2016).

At the late metatrochophore stage, *Avi-en* metameric domains were found within the territories of two adjacent segments (Fig. 2F), likely indicating the abutting or even overlapping expression of *Avi-wnt1* (Fig. 5M) and *Avi-en* along the posterior segment border. Similar relationships were observed during postlarval segment development: initially, *Avi-en* was expressed only at the anterior border of the segment primordium. Later, its expression expanded across the furrow to reach the posterior border of the adjacent anterior segment (Fig. 2N), while *Avi-wnt1* remained confined to the posterior border of the segment (Fig. 5S). During both trochophore and metatrochophore stages of larval development, adjacent expression of *Avi-wnt1* and *Avi-en* was observed along the segment-pygidium border (Fig. 3, 6). Notably, the *Avi-wnt1* stripe at the posterior border of the third larval segment was maintained until the *Avi-en* domain in the growth zone began to expand and thicken in early nectochaetes (180 h.p.f.). This indicates that the stable posterior ring of *Avi-en* expression is periodically flanked by an anterior band of *Avi-wnt1*, a relationship that holds true for both larval and postlarval segment formation. These findings strongly support the existing model, proposed for caudal regeneration in *Perinereis nuntia*, wherein Wnt signaling induces the formation of new segmental boundaries (Niwa et al., 2013).

The posterior medial domain of *Avi-wnt1* was confined to the proctodaeum (Fig. 5, 7). Starting at the metatrochophore stage, this domain developed into two parts: a dorsal and a ventral region (Fig. 5E). By the late metatrochophore stage, *Avi-wnt1* expression became detectable internally, within the hindgut anlage (Fig. 5K), likely due to invagination during hindgut morphogenesis, as previously noted (Kulakova et al., 2008).

Significant changes in *Avi-en* expression occurred in the most posterior domain during the formation of the first postlarval segment. At the late metatrochophore stage, *Avi-en* was expressed in a circular arrangement of superficial (ectodermal) and some inner (mesodermal) cells. Subsequently, at the early nectochaete stage, this domain expanded along the AP axis, forming two rows of superficial cells (Fig. 2G, 8). These rows in the pygidium eventually separated due to cell proliferation, as demonstrated by the colocalization of *Avi-en* expression signal with EdU labeling (Fig. 4). These processes reflect the sequential steps involved in the formation of the first postlarval segment (Kairov and Kozin, 2023). During subsequent postlarval growth, the emergence of new expression stripes of *Avi-en* and *Avi-wnt1* consistently preceded the formation of the segment rudiment (Fig. 8).

Thus, the expression of *Avi-wnt1* and *Avi-en* accompanied the development of both larval and postlarval segments. In both cases, the metameric *Avi-en* pattern emerged as a result of the subdivision of an initially continuous expression domain. In contrast, *Avi-wnt1* expression appeared later than *Avi-en* at the segment boundaries and exhibited a metameric nature from its onset. The mechanisms inducing *Avi-en* and *Avi-wnt1* metameric expression in the hyposphere remain the most challenging question.

### Evolutionary conserved and derived aspects of *Avi-en* and *Avi-wnt1* expression

The dynamics of *Avi-en* and *Avi-wnt1* expression reveal numerous developmental traits in *A. virens*, some unique to nereidid polychaetes and others conserved across annelids. In the embryos of *A. virens* and *P. dumerilii*, *en* and *wnt1* transcripts appear early in development, specifically during the protrochophore stage. By the mid-trochophore stage, the expression domains are clearly associated with the pygidium and future segments (Prud’homme et al., 2003; Steinmetz et al., 2011). The dynamics of *en* expression in nereidids resemble those in another polychaete with lecithotrophic larvae, *Capitella teleta*. In all these species, *engrailed* mRNAs initially appear at the lateral sides of the hyposphere, with expression domains subsequently expanding to the ventral and dorsal sides (Seaver and Kaneshige, 2006). However, in *C. teleta*, *engrailed* expression begins only after the differentiation of ciliary bands, whereas in *A. virens*, it occurs prior to ciliation. This difference can be interpreted as heterochrony, implying accelerated pattern formation in nereidids. In the basal polychaete *Chaetopterus* sp., *engrailed* expression begins in the L4 larva at the lateral sides of the body. This pattern later resolves into metameric domains and expands dorsally (Seaver et al., 2001), but only after the formation of morphological segment boundaries. Another common feature across these species is the detection of *engrailed* mRNAs in the mesoderm of the prospective growth zone. In *A. virens* and other studied polychaetes (*C. teleta*, *Chaetopterus* sp., and *Hydroides elegans*), *engrailed* expression first appears in superficial cells, followed by expression in deeper mesodermal cells. This pattern may arise from various processes, such as the hypothetical migration of cells from the surface to the interior (Seaver and Kaneshige, 2006) or as a result of inductive interactions. This feature of expression appears to be conserved among annelids but is absent in leeches. In the latter, *engrailed* expression is detected in the mesodermal descendants of teloblasts (Lans et al., 1993), but not in the teloblasts themselves, which together constitute the embryonic growth zone.

In the sedentary polychaete with planktotrophic larvae *H. elegans*, *engrailed* mRNA is detected starting only from the trochophore stage, first in the ectoderm, and then metamerically in deep cells (future bristle-bearing sacs) (Seaver and Kaneshige, 2006). By the time of morphological segmentation, the signal is observed in the mesoderm. A comparison of the dynamics of *engrailed* expression shows that in nereidids and *C. teleta* (polychaetes with lecithotrophic larvae), expression occurs earlier than in species with planktotrophic larvae. Moreover, in the echiurid *Urechis unicinctus*, transcripts of *engrailed* and other genes associated with segmentation are up-regulated quite early in development (Hou et al., 2019). All of this confirms the results of the recent study, which proposes that species with lecithotrophic larvae develop a trunk earlier than species with planktotrophic larvae (Martín-Zamora et al., 2023).

*Engrailed* genes are known to participate in the specification of many serial structures, such as CNS ganglia, nephridia, and chaetal sacs. All studied annelids showed *engrailed* expression in the nervous system (Patel et al., 1989; Bely and Wray, 2001; Seaver et al., 2001; Prud’homme et al., 2003; Seaver and Kaneshige, 2006; Kairov and Kozin, 2023). Moreover, neural expression is identified in other bilaterian animals, such as vertebrates, arthropods, and echinoderms (Condron et al., 1994; Byrne et al., 2005; Omi and Nakamura, 2015). These data suggest that the hypothesis proposed for arthropods about the ancestral expression of *engrailed* in the metameric elements of the nervous system and subsequent co-option in the process of segment establishment and patterning may also apply to annelids (Patel et al., 1989; Chipman, 2010).

The presence of *wnt1* at the posterior end of the body is a conserved feature of bilaterians (Martin and Kimelman, 2009; Petersen and Reddien, 2009; Loh et al., 2016). The corresponding *Avi-wnt1* expression domain is the earliest one in *A. virens*. Among annelids, *wnt1* expression in the hindgut anlage was characterized in *C. teleta* and the leech *Helobdella robusta* (Seaver and Kaneshige, 2006; Cho et al., 2010). There is metameric *wnt1* expression in their trunk, ectodermal in the leech, and mesodermal in *C. teleta*. Other *wnt* paralogs are expressed metamerically too, forming ectodermal stripes (e.g., *wnt16a* and *wnt16b* in *H. robusta*). Nereidid polychaetes are also characterized by metameric *wnt* expression both in larval and postlarval development (Prud’homme et al., 2003; Janssen et al., 2010; Pruitt et al., 2014; Kozin et al., 2019). Widely distributed metameric Wnt activity in annelids suggests that Wnt signaling was involved in segmentation and body patterning in the annelid ancestor.

While our study focuses on the expression of *Avi-en* and *Avi-wnt1*, it is important to consider that segmentation likely involves a more complex interplay of multiple Wnt ligands. In the annelid *P. dumerilii*, several Wnt genes, including *wnt1, wnt10, wnt11,* and *wnt16*, are expressed in overlapping domains at segment boundaries, suggesting potential redundancy or combinatorial action (Janssen et al., 2010). A similar diversity of Wnt paralogs has also been described for *A. virens*, corresponding to the repertoire found in *P. dumerilii* (Kozin et al., 2019). Furthermore, comprehensive expression analysis in the leech *Helobdella robusta* revealed that duplicated Wnt paralogs (e.g., *wnt5, wnt11, wnt16*) are often expressed in distinct yet partially overlapping patterns, indicating subfunctionalization and the formation of complex “Wnt landscapes” (Cho et al., 2010). Although not detected in our assays, the potential co-expression of other Wnt paralogs could influence the expression dynamics of *Avi-en*. Therefore, the complementary patterns we describe for *Avi-en* and *Avi-wnt1* may be part of a more extensive regulatory network, whose full complexity remains to be elucidated.

Comparison of segmentation gene activity in annelids and arthropods provides rich material for evolutionary interpretations. Despite the similarities in *wnt1* and *engrailed* patterns between nereidids and other annelids, only nereidids have a complementary and striped arrangement of *wnt1* and *engrailed* domains of expression resembling arthropods. Virtually all arthropods and related to them onychophorans and tardigrades have metameric distribution of *wnt1* and *engrailed* mRNAs during parasegment development (Gabriel and Goldstein, 2007; Eriksson et al., 2009; Williams and Nagy, 2017). But the most studied annelids do not exhibit such a pattern. Leeches have metameric *engrailed* expression, but ablation of *engrailed*-positive cells doesn’t affect segmentation (Seaver and Shankland, 2001). Therefore, nereidids appear to have gained a new mechanism of segment patterning similar to arthropods but lacking in other annelids. Furthermore, the temporal sequence of the appearance of the *wnt1* and *engrailed* metameric bands is similar — in arthropods, during terminal growth, the *engrailed* band appears before the *wnt1* band (Chesebro et al., 2013; Lim and Choe, 2020). In *A. virens*, the metameric expression of *engrailed* appears much earlier than *wnt1* and, unlike in arthropods, is present at both the anterior and posterior parts of the segment during larval and postlarval development (Fig. 2F, N). This indicates a fundamentally different interaction between *engrailed* and Wnt signaling in the segment patterning of *A. virens* compared to arthropods.

*Engrailed* activity at the borders of segments and the pygidium in nereidids supports the hypothesis of the ancestral role of *engrailed* in boundary formation (Vellutini and Hejnol, 2016). Our data align with the evolutionary scenario in which the expression of *engrailed* was not initially associated with segmentation, and *engrailed* was likely co-opted into segmentation independently in different phylogenetic lineages. The exact timing of this co-option in nereidids remains unclear, as this expression pattern has been revealed only in this family. Gene expression studies in other annelid families are essential to clarify this issue.

The expression of *engrailed* in the growth zone is characteristic of many studied annelids (Bely and Wray, 2001; Prud’homme et al., 2003; Seaver and Kaneshige, 2006). In *Chaetopterus* sp., expression of *engrailed* posterior to the C2 segment can be interpreted as expression in the growth zone (Seaver et al., 2001). In both *C. teleta* and *Chaetopterus* sp., *engrailed* mRNA is detected in the growth zone only in pre-metamorphic larvae, whereas in *A. virens*, the most posterior band of *engrailed* expression is evident as early as the trochophore stage. Spatially, this domain corresponds to the future expression region of multipotency markers (Rebscher et al., 2007; Kozin and Kostyuchenko, 2015; Kostyuchenko, 2022). Based on this, we hypothesize that in *A. virens* (and other nereidids), there is an early subdivision of the hyposphere into the segmental part and the future pygidium.

To summarize, *A. virens* exhibits both conserved features in the expression of *engrailed* and *wnt1* (such as the presence of *engrailed* transcripts in the elements of the nervous system, and in the ectoderm and mesoderm of the growth zone, as well as *wnt1* expression in the hindgut and metameric domains), as well as traits unique to nereidid polychaetes. These include the early induction of *engrailed* expression in the growth zone and segmental rudiments, as well as the complementary arrangement of the *engrailed* and *wnt1* expression domains.

### Relationships between larval and postlarval segmentation

The revealed differences in *engrailed* expression during the larval period across various annelids prompt us to refer to Ivanov’s theory on the primary heteronomy of segments (Iwanoff, 1928; Schroeder and Hermans, 1975; Ivanova-Kazas, 1978; Giangrande and Gambi, 1998). The similar pattern of *engrailed* expression in polychaete larvae—characterized by the subdivision of a continuous longitudinal domain into distinct metameric spots, expanding from the lateral sides to the dorsal and ventral sides (Fig. 7)—may indicate a more conservative mode of larval segment patterning. However, no such similarity was observed in the molecular patterning of postlarval segments, which may support the idea that larval segment development represents an ancestral characteristic. The exact ancestral mechanisms involved in segment specification in annelids remain controversial. In clitellates (oligochaetes and leeches), germ-band cells acquire their identity upon the division of teloblasts. These nascent primary blast cells proliferate stereotypically and assume specific positions within a segment, a process known as lineage-driven segmentation (Weisblat and Kuo, 2014; Weisblat and Winchell, 2020). Whether this process applies to non-clitellate annelids remains uncertain. In *P. dumerilii* early development, teloblastic activity resembling that of clitellate annelids was observed in mesodermal bands, but not in the ectoderm (Fischer and Arendt, 2013; Özpolat et al., 2017). In *C. teleta* and *H. elegans*, no teloblasts have been described in either the ectoderm or mesoderm (Seaver et al., 2005; Irvine and Seaver, 2006). However, in *C. teleta*, *nanos* expression was detected in a distinct ring of ectodermal cells anterior to the telotroch, which may suggest the presence of ectodermal teloblasts (Dill and Seaver, 2008). Morphologically distinct teloblasts have been described in the larva of *Scoloplos armiger* (Anderson, 1959), but there is no molecular or functional data to confirm their identity.

**Fig. 7.**
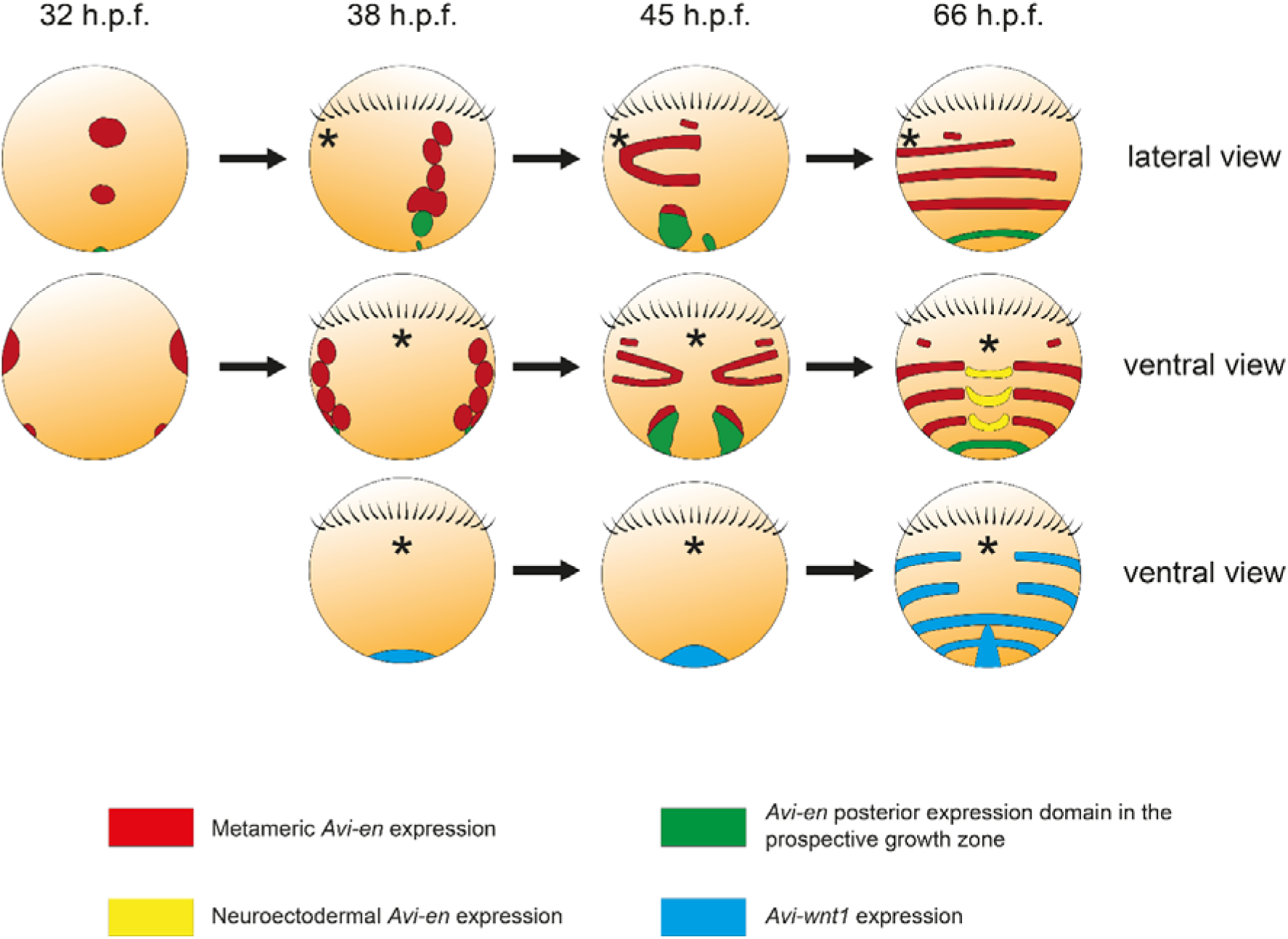
Schematic representation of *Avi-en* and *Avi-wnt1* expression, which patterns larval segments during early stages of *A. virens* development. Anterior is to the top. Black asterisks — stomodeum, black strokes — prototroch. Upper row — *Avi-en* expression, lateral view; middle row — *Avi-en* expression, ventral view, lower row — *Avi-wnt1* expression, ventral view.

**Fig. 8.**
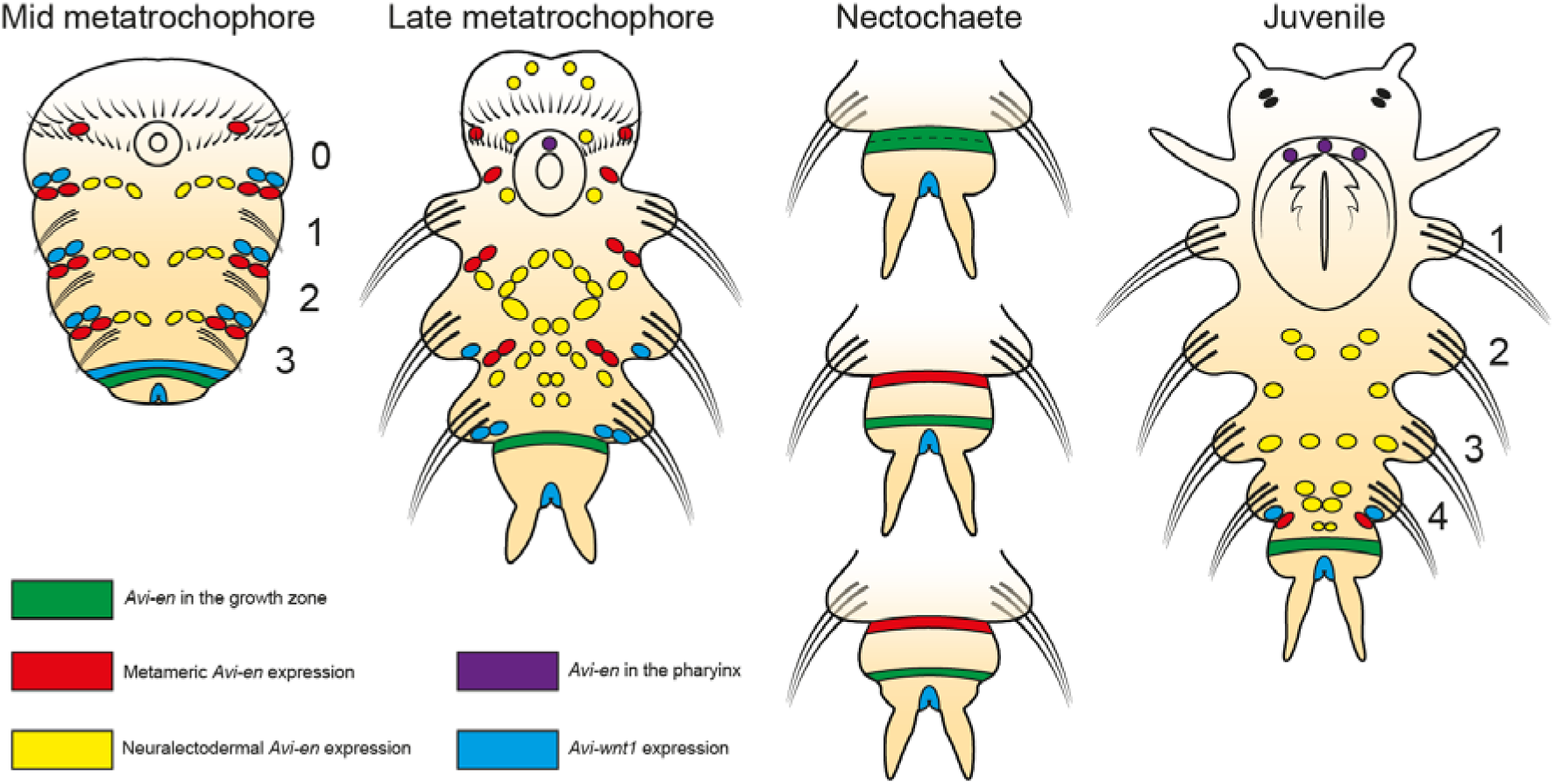
Schematic representation of *Avi-en* and *Avi-wnt1* expression during formation of the first postlarval segment in *A. virens*. Anterior is to the top. Developmental stages are indicated above. Each segment is numbered with an Arabic numeral.

The question of the origin of the cellular material for segments is crucial in the evolutionary analysis of segmentation (Minelli, 2005). At least in nereidids, it can be confidently stated that larval segments arise from widespread proliferation and migration of cells in the hyposphere, leading to its convergent extension (Denes et al., 2007; Steinmetz et al., 2007; Demilly et al., 2013). These processes could explain the separation of the originally united lateral longitudinal domains of *engrailed* expression (Fig. 1, 7). In contrast, postlarval segments derive from the proliferation of subterminal growth zone cells and the cellular material of each new segment anlage. In clitellate annelids, these differences do not exist, as all segments are formed by local cell proliferation. By analogy with clitellates, some authors have suggested that the source of larval segments in non-clitellate annelids is the growth zone of the trochophore (Anderson, 1973; Shimizu and Nakamoto, 2001), but this speculation is inconsistent with modern observations.

Inductive interactions between the ectoderm and mesoderm also play a critical role in segment specification. In clitellate embryos, the mesoderm governs segment morphogenesis and the axial identity of the ectoderm (Blair, 1982; Shimizu et al., 2001). Whether such interactions are present in non-clitellate annelids remains unknown, but the metameric pattern of *Avi-twist* expression in mesodermal bands (Kozin et al., 2016) forms even earlier than *Avi-en* expression in the ectoderm (Fig. 1E). This is strikingly different from arthropods, where the ectoderm determines mesoderm segmentation (Azpiazu et al., 1996; Hannibal et al., 2012). To assess the autonomy of specification, it would be necessary to limit external signaling influences on the region undergoing segmentation, a task that could be accomplished through explantation or complete inhibition of signaling interactions.

Several studies on the inhibition of intracellular processes have revealed segmentation abnormalities. Notably, interesting results were obtained in *A. virens* embryos treated with antibiotics (actinomycin D and sibiromycin) during gastrulation and trochophore formation. These treatments led to the formation of unsegmented larvae instead of segmented metatrochophores (Dondua, 1975). This phenotype appears incompatible with a lineage-driven mode and may thus support the inductive specification of larval segments. In line with this, inhibition of Hedgehog signaling in *P. dumerilii* (Dray et al., 2010) also led to a reduction in ectodermal striped expression of *hedgehog*, *engrailed*, *wnt1*, and *lbx*. Notably, the reduction in segmental grooves in larval segments was more pronounced than in postlarval segments. The authors concluded that Hedgehog is not necessary to establish early segment patterns but is required to maintain them. Additionally, segmentation of regenerating posterior body ends was impaired due to modulation of Wnt (Niwa et al., 2013; Ribeiro and Aguado, 2021) and FGF signaling pathways (Shalaeva et al., 2021; Shalaeva and Kozin, 2025). In contrast, Delta/Notch signaling does not appear to be involved in segmental pattern formation during either the larval or postlarval periods (Thamm and Seaver, 2008; Rivera and Weisblat, 2009; Gazave et al., 2017).

### Sequential segmentation and the growth zone organization

The organization of the growth zone (also referred to as the segment addition zone, or SAZ, in arthropods) is central to understanding the origin of segmentation in bilaterians. Among annelids, the growth zone has been well studied in clitellates, particularly in leeches, where it consists of several large stem cells (teloblasts) that give rise to all body segments (Weisblat and Kuo, 2014; Zattara and Weisblat, 2020). However, based on the limited distribution of teloblasts and the existing hypotheses regarding the evolution of annelid ontogeny, we propose that a teloblastic SAZ is not the ancestral trait of annelids (Scholtz, 2002; Minelli, 2005; Kuo, 2017). Instead, it is likely a secondary gain, meaning clitellates are not the most appropriate model for comparative analysis.

Currently, we lack a clear understanding of how the growth zone functions in non-clitellate annelids. Most of the available data comes from nereidid polychaetes. Studies on *P. dumerilii* have proposed the existence of a teloblastic growth zone, although teloblasts do not differ in size from other cells (Gazave et al., 2013; Balavoine, 2014). However, there are no signs of stereotypical division patterns in this zone, except for the longitudinal orientation of mitoses in clusters of ectodermal growth zone cells during regeneration in *Perinereis nuntia* (Niwa et al., 2013). The diffuse model of SAZ, characteristic of arthropods and vertebrates, which involves periodic gene expression (e.g., *hairy* in vertebrates or pair-rule genes in arthropods) (Bénazéraf and Pourquié, 2013; Williams and Nagy, 2017), is not applicable to nereidids. The diffuse SAZ model designates a wide area of unsegmented tissue along the AP axis, where the posterior shift of the wavefront results in a gradual decrease in differentiation potency, leading to segment specification. This pattern is referred to as boundary-driven segmentation (Weisblat and Kuo, 2014). However, no oscillations in gene expression have been detected in annelids to date.

One of the most critical mechanisms for segment addition is cell proliferation. In arthropods and vertebrate embryos, discrete regions of cell proliferation are observed (Venters et al., 2008; Nagy and Williams, 2020). In particular, cell divisions in short germ-band arthropods occur in the posterior part of the growth zone and in the last specified metamer (Fig. 9A-C). Annelids, in contrast, exhibit ubiquitous proliferation in the hyposphere of the trochophore (Demilly et al., 2013) and in a continuous, posterior subterminal region, extending from the compact growth zone to the newly formed segments (Fig. 4, 9A’-C’). This pattern of cell division appears to be unique to the segment-producing tissues in annelids, both clitellate (Bissen and Weisblat, 1989) and non-clitellate. Consequently, the discrete pattern of cell divisions in a diffuse SAZ is incompatible with the mechanisms forming both larval and postlarval segments in annelids.

**Fig. 9.**
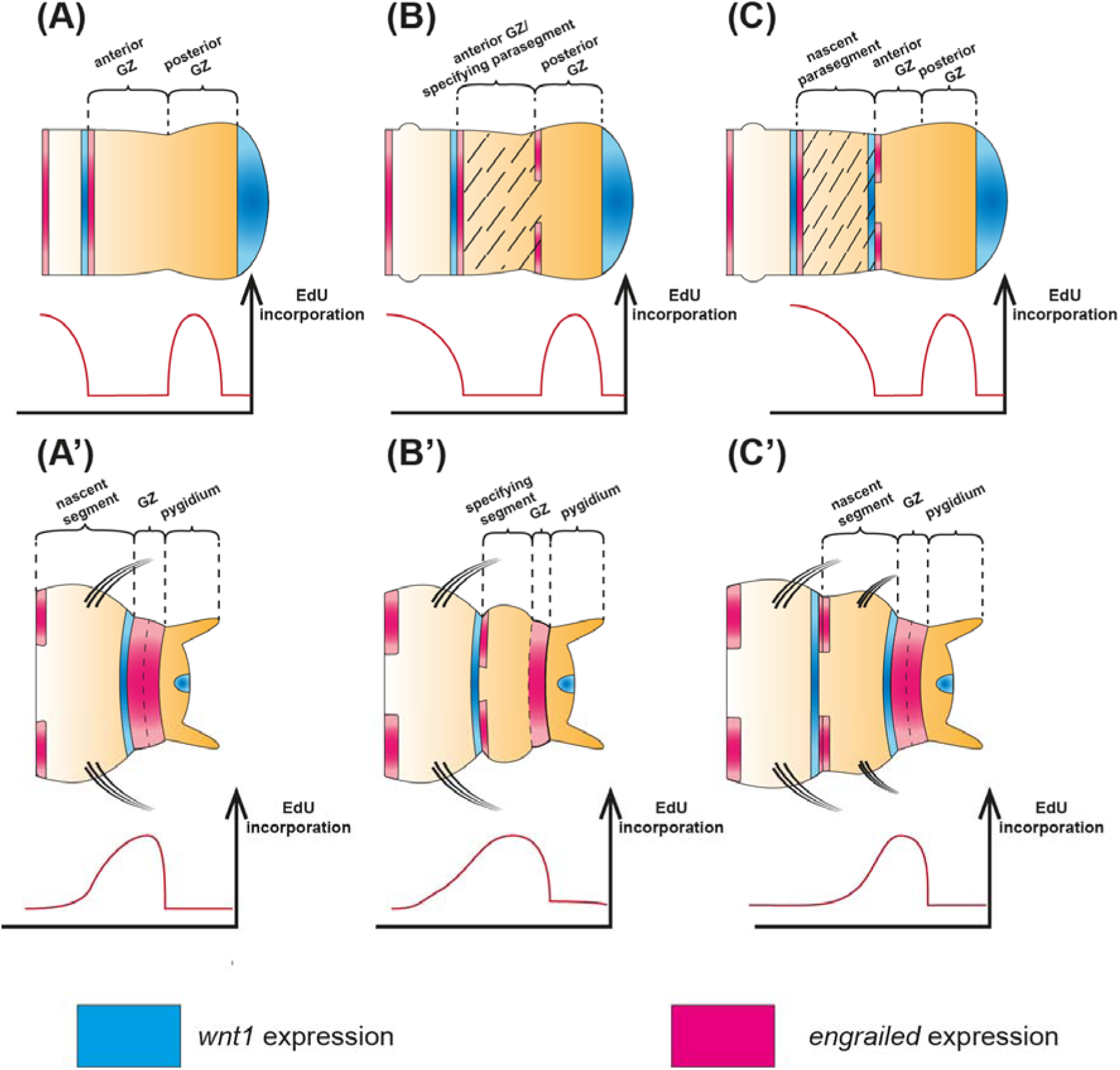
Comparison of sequential segment formation between Arthropods with the diffuse SAZ (upper row) and *A. virens* (lower row). Dashed area — specifying metamer (parasegment). Blue areas — *wg*/*wnt1* expression, red areas — *engrailed* expression. Charts below each diagram represent EdU incorporation level along the AP axis (Nagy and Williams, 2020). (A) Cell accumulation in the anterior growth zone. (B) Gene expression oscillations (pair-rule genes or Notch-signaling components) promote parasegment specification. (C) Determination of parasegment borders (segment polarity genes expression). (A’) Expansion of the posterior *Av-en* domain indicates definition of the new segment’s anterior border. (B’) Cell divisions gradually produce more posterior territories of the specifying segment. (C’) *Avi-wnt1* expression in front of segment-pygidium furrow indicates determination of the whole new segment anlage.

Our results suggest several key features of sequential segment addition in nereidid annelids (Fig. 9). We distinguish three stages of postlarval segment development: elongation of the AP axis, metamer specification and separation (which ultimately determines segmental borders). Unlike arthropods, nereidids exhibit *engrailed* expression in the growth zone (Fig. 9A’), which resolves into two rows by the nectochaete stage (Fig. 9B’). According to our data and previous studies (Niwa et al., 2013), *wnt1* expression at the posterior border of the developing metamer represents the final step in the segmentation program (Fig. 9C’). Thus, axial elongation and segment specification occur simultaneously in nereidids. This contrasts with parasegment formation in arthropods, where these processes are strictly sequential (Williams and Nagy, 2017). These differences cast doubt on the homology of segments among bilaterians, as proposed by numerous authors (Scholtz, 2002; Balavoine and Adoutte, 2003; Prud’homme et al., 2003; de Rosa et al., 2005; Saudemont et al., 2008; Couso, 2009; Balavoine, 2014; Malakhov et al., 2019; Malakhov and Gantsevich, 2022; Shcherbakov, 2023). Our data suggest that the involvement of *engrailed* and *wnt1* in segmentation may be independent in annelids and other taxa (Seaver, 2003, 2022; Chipman, 2010; Ferrier, 2012; Bleidorn et al., 2015; Zattara and Weisblat, 2020; Kairov and Kozin, 2023), which is also supported by *engrailed* patterns in diverse metazoans (Vellutini and Hejnol, 2016). This scenario is further reinforced by the fact that segmental grooves in arthropods are formed posteriorly from the *engrailed* expression band, while in nereidid annelids, they are formed anteriorly from the *engrailed* band and posteriorly from the *wnt1* band (Fig. 9). Annelid segments have been homologized with parasegments in arthropods (Prud’homme et al., 2003; Janssen et al., 2010), since parasegmental furrows are formed between the *wnt1* and *engrailed* expression bands. However, these parasegmental furrows may have convergent evolutionary origins and are not considered to be a conserved feature across arthropods (Janssen et al., 2022).

Thus, the model of sequential segmentation in nereidids involves the gradual formation of a segment along the AP axis (Kairov and Kozin, 2023). The specification of a new segment begins as *engrailed* expression in the growth zone expands through cell divisions. This widened ring of *engrailed-*positive cells presumably give rise to the anterior border of the new metamer and to the maintaining growth zone posteriorly. Subsequent proliferation splits the *engrailed* expression bands, which is morphologically manifested by pygidium growth. We hypothesize that descendant cells leaving the growth zone downregulate *engrailed* and exhibit higher proliferation rates than the cells retained in the growth zone itself, consistent with BrdU incorporation data from *P. dumerilii* (de Rosa et al., 2005). Segment specification is complete when the *wnt1* expression band appears, marking the posterior boundary of the segment. To further validate this model, future studies should employ advanced cell proliferation assays, such as pulse-chase EdU labeling and the application of additional markers.

The previous model of gradual segment specification was first proposed for regenerating *P. nuntia* (Niwa et al., 2013). In this model, the growth zone is not a population of bona fide multipotent stem cells but rather a region where the anterior-most pygidial cells undergo reiterative transdifferentiation. However, this does not align with the consistent *engrailed* expression at the anterior border of the pygidium, which is observed not only in nereidids but also in *C. teleta* and *Pristina leidyi* (Bely and Wray, 2001; Seaver and Kaneshige, 2006). The model proposed by Niwa and colleagues (2013) suggests that Wnt signaling induces segment precursors, as Wnt hyperactivation alters the size and number of emerging segments, which is proved in *P. nuntia* and *Sillys malaquini* (Niwa et al., 2013; Ribeiro and Aguado, 2021). We therefore propose that Wnt signaling regulates the transition from segment elongation to the initiation of a new segment, a process characterized by the appearance of adjacent *wnt1* and *engrailed* expression domains at the anterior pygidial border. The *Avi-wnt1* expression patterns in *A. virens* are consistent with the hypothesis that Wnt signaling plays a crucial role in the patterning of both larval and postlarval segments.

Further interpretation of segmentation traits in nereidids leads us to hypothesize that *Avi-wnt1* expression in the proctodaeum/hindgut is required for the induction and maintenance of growth zone function, whereas Wnt activity in specified segments is involved in their polarization and in regulating the pace of segment addition, as reflected in the size of the newly forming segment. The terminal posterior Wnt signaling center is highly conserved across bilaterians (Martin and Kimelman, 2009; Petersen and Reddien, 2009; Loh et al., 2016), and it plays a critical role in axial elongation in both segmented (Shimizu et al., 2005; Bolognesi et al., 2008; Dunty et al., 2008; Chesebro et al., 2013) and non-segmented animals (Fritzenwanker et al., 2019). Posterior Wnt activity has been confirmed in all annelids studied (Seaver et al., 2001; Prud’homme et al., 2003; Seaver and Kaneshige, 2006; Cho et al., 2010; Janssen et al., 2010; Nyberg et al., 2012; Niwa et al., 2013; Pruitt et al., 2014; Kozin et al., 2019; Ribeiro and Aguado, 2021), whereas Wnt expression in stripes along segmental borders has only been reported in nereidids and *C. teleta* (Cho et al., 2010). To clarify how segmented body plans evolved within annelids, future studies should primarily focus on Wnt-mediated segmentation mechanisms across different phylogenetic lineages of non-clitellate annelids.

## Acknowledgements

This research was funded by the RSF grant 23-74-10046, https://rscf.ru/en/project/23-74-10046/. We are grateful to the research resource centers “Microscopy and Microanalysis”, “Molecular and Cell Technologies”, “Chromas”, “Culture Collection of Microorganisms”, and the Marine Biological Station of St. Petersburg State University for their technical support. We thank Roman P. Kostyuchenko for plasmids with fragments of the cloned cDNA of *Avi-wnt1* and *Avi-en*.

## Notes

### Competing Interest Statement

The authors have declared no competing interest.

### Summary of Updates

We performed new experiments to clarify en and wnt1 expression patterns, the results of which are now included as new Fig. 6. The staining patterns from the mixed probes are broader than those from individual probes, indicating that Avi-en and Avi-wnt1 are expressed in adjacent but largely non-overlapping cell populations. This suggests a signaling relationship rather than co-expression. Accordingly, we corrected representation of expression patterns on figures 8 and 9.

